# Refinement of intraperitoneal injection of sodium pentobarbital for euthanasia in laboratory rats (*Rattus norvegicus*)

**DOI:** 10.1101/081414

**Authors:** K Zatroch, CG Knight, JN Reimer, DSJ Pang

## Abstract

**Background:** The Canadian Council on Animal Care and American Veterinary Medical Association classify intraperitoneal (IP) pentobarbital as an acceptable euthanasia method in rats. However, federal guidelines do not exist for a recommended dose or volume and IP euthanasia has been described as unreliable, with misinjections leading to variable success in ensuring a timely death. The aims of this study were to assess and improve efficacy and consistency of IP euthanasia.

**Methods:** In a randomized, blinded study, 51 adult female Sprague-Dawley rats (170-495 g) received one of four treatments: low-dose low-volume (LL) IP pentobarbital (n = 13, 200 mg/kg pentobarbital), low-dose high-volume (LH) IP pentobarbital (n = 14, 200 mg/kg diluted 1:3 with phosphate buffered saline), high-dose high-volume (HH, n = 14, 800 mg/kg pentobarbital), or saline. Times to loss of righting reflex (LORR) and cessation of heartbeat (CHB) were recorded. To identify misinjections, necropsy examinations were performed on all rats. Video recordings of LL and HH groups were analyzed for pain-associated behaviors. Between-group comparisons were performed with 1-way ANOVA and Games-Howell post hoc tests. Variability for CHB was assessed by coefficient of variation (CV) calculation.

**Results:** The fastest euthanasia method (CHB) was HH (283.7 ± 38 s), compared with LL (485.8 ± 140.7 s, p = 0.002) and LH (347.7 ± 72.0 s, p = 0.039). Values for CV were: HH, 13.4%; LH, 20.7%; LL, 29.0%. LORR time was longest in LL (139.5 ± 29.6 s), compared with HH (111.6 ± 19.7 s, p = 0.046) and LH (104.2 ± 19.3 s, p = 0.01). Misinjections occurred in 15.7% (8/51) of euthanasia attempts. Pain-associated behavior incidence ranged from 36% (LL) to 46% (HH).

**Conclusion:** These data illustrate refinement of this euthanasia technique. Both dose and volume contribute to speed of death with IP pentobarbital and an increase in volume alone does not significantly reduce variability. The proportion of misinjections was similar to that of previous studies.

**Abbreviations:**

LL: low-dose low-volume
LH: low-dose high-volume
HH: high-dose high-volume
LORR: loss of righting reflex
CHB: cessation of heartbeat
GIT: gastrointestinal tract
CV: coefficient of variation

## sIntroduction

Over 2 million rats are used in biomedical research in Canada and the European Union annually [1, 2]. The overwhelming majority of laboratory studies employing rodents end with killing the animals upon completion of the study or if a humane endpoint has been reached. While this is a reality of research, efforts to refine killing methods, to achieve “euthanasia”, for rats and other laboratory animals are ongoing, as reflected in recent updates to the Canadian Council on Animal Care (CCAC) and American Veterinary Medical Association (AVMA) euthanasia guidelines [3, 4]. Goals for successful euthanasia include techniques requiring minimal restraint, simplicity of administration, and a swift, painless death [5].

A commonly employed technique for euthanasia of laboratory rats is an overdose of carbon dioxide. However, current behavioral and physiologic evidence suggests that this method is aversive and may be painful [6-15]. As a result, the AVMA and CCAC have reclassified killing with carbon dioxide as “conditionally acceptable” [4] and “acceptable with conditions” [3].

In contrast, an acceptable method and preferred alternative to carbon dioxide is overdose with a barbiturate such as sodium pentobarbital (PB). An intraperitoneal (IP) route of injection is acceptable when intravenous injection cannot be performed or is impractical [3, 4]. Current guidelines do not indicate a specific dose of sodium pentobarbital for euthanasia, although 200 mg/kg or 3 times the anesthetic dose have been suggested [5]. There are several potential drawbacks associated with IP PB injection, including misinjection, variability in effect and pain [7, 16–22].

An important factor contributing to variability of drug effect (speed of onset and success) is misinjection, with deposition of injectate into intra-abdominal fat, abdominal viscera or the subcutaneous space. In the case of IP pentobarbital for euthanasia this results in a delayed time to death or even failure to cause loss of consciousness. Attempts to reduce variability with a two-person injection technique (one to restrain, one to inject) have had variable success, with reported proportions of misinjections ranging from 6 to 20% [18–20].

Pain, inferred from behavioral observations, necropsy findings and biomarkers, has also been cited as a potential impediment to achieving the principle of euthanasia.

Specifically, exhibition of writhing (defined as the contraction of the abdomen and extension of the hind legs), grossly visible inflammation of abdominal viscera at necropsy and a measurable increase in spinal cord cFos have been reported following IP injection of pentobarbital [16, 17, 21, 22].

The primary aim of this study was to assess the impact of varying the dose and volume of sodium pentobarbital injected into the intraperitoneal cavity on time to death and consistency of the killing process. Secondary aims were identification of misinjections by necropsy and the quantification of writhing behavior in response to IP PB. We hypothesized that speed and consistency of IP euthanasia would be improved by using a higher dose and higher volume.

## Methods

### Ethics Statement

The animal care and use protocol was approved by the Health Sciences Animal Care Committee of the University of Calgary (AC11-0044), in accordance with the guidelines of the CCAC.

### Study Design

51 adult female Sprague-Dawley rats (170-495 g), sourced as surplus breeding stock, were included in the study. A sample size of approximately 13 animals, to achieve 80% power with an alpha of 0.05 (with an anticipated 20% misinjection rate) with an effect size of 1.5, was determined from pilot data. All animals remained in paired housing until the time of trial and were not handled prior to the study. Housing consisted of standard micro-filter cages (47 X 25 X 21 cm) with wood shavings and shredded paper bedding and a plastic tube for enrichment. A 12-12 hour lights on-off cycle (lights on at 0700) was maintained in an environmentally controlled room (23˚C, 22% humidity). All experiments were performed during the light period (0730-1800).

Animals were randomly assigned to one of four treatment groups for IP injection. A low-dose low-volume (LL, n = 13) group received 200 mg/kg sodium pentobarbital (Euthanyl, 240 mg/ml, Bimeda-MTC Animal Health Inc., Cambridge, ON, Canada). A low-dose high-volume group (LH, n = 14) received 200 mg/kg sodium pentobarbital diluted 1:3 with phosphate-buffered saline (PBS). A high-dose high-volume (HH, n = 14) group received 800 mg/kg sodium pentobarbital. A control group (n = 10) received 1 ml of PBS. Each treatment was placed in a 1 ml (LL and control groups) or 3 ml (LH and HH groups) syringe as dictated by the volume of injectate. A new 25 G 5/8” hypodermic needle was attached to each syringe for injection. Blue food coloring (0.01 mL, Club House, Burlington, Ontario) was added to each treatment to facilitate visualization of injectate during necropsy examination.

At the beginning of each trial, a single rat was removed from the housing unit and placed in a Plexiglas chamber (L X W X H: 27.5 X 14.5 X 20.5 cm). Two video cameras (Panasonic HC-V720P/PC, Panasonic Canada Inc., Mississauga, ON, Canada) were placed along the long and short axes of the chamber. Prior to each injection, baseline video of the rat was recorded for 10 minutes. Treatments were prepared in a separate room during baseline video recording. Individuals performing the IP injections and behavioral analyses were blinded to treatment.

Following baseline video, each rat was removed from the box and restrained for a two-person injection technique. Rats were held in dorsal recumbency at an approximately 30-degree angle (head lowermost). The holder (DP) supported each rat and restrained the left pelvic limb. The individual administering each injection (KZ) restrained each rat’s right pelvic limb, injecting with the right (dominant) hand (Fig. 1). Each injection was performed in the right caudal quadrant of the abdomen at the level of the coxofemoral joint and approximately 5 mm to the right of midline. The needle was directed cranially at a 45-degree angle to the body wall.

**Fig 1:**
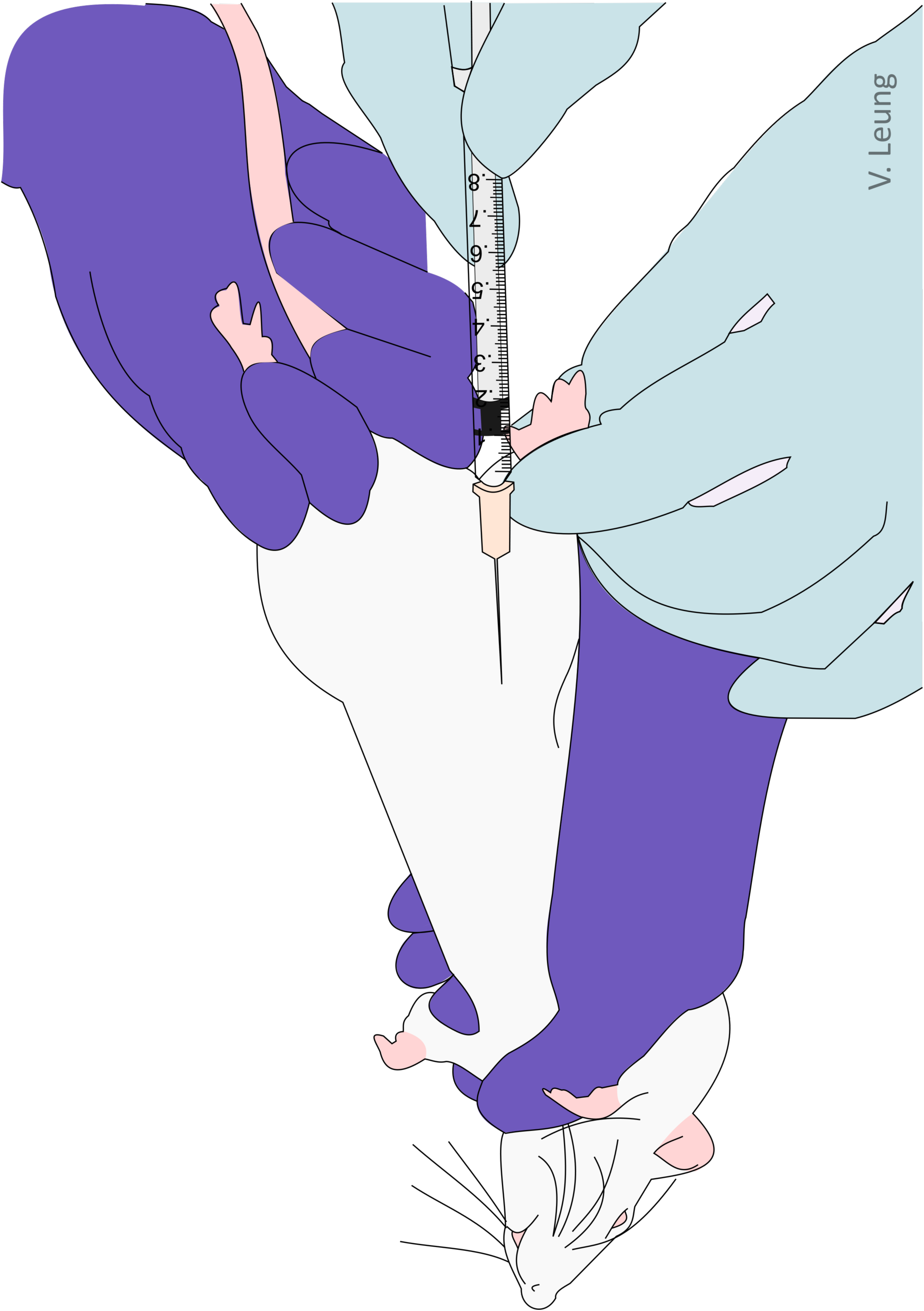
A cartoon showing the two-person injection technique used in the study, with one person holding the rat in dorsal recumbency (head down) and the second person gently restraining the right pelvic limb to facilitate intraperitoneal injection in to the right caudal quadrant.

Immediately following completion of injection, each rat was returned to the observation chamber. A single blinded observer (KZ) monitored for signs of ataxia (stumbling, falling, crossing feet) following injection. If signs of ataxia were noted, an attempt was made to place the rat in dorsal recumbency to evaluate for a loss of righting reflex (LORR), a surrogate for loss of consciousness [7, 23]. LORR was confirmed if the rat remained in dorsal recumbency for ten seconds. Failure of LORR was established if the rat resisted initial placement on its back or was able to right itself within ten seconds. In cases of initial LORR failure, the test was repeated every 30 seconds until LORR occurred. Following LORR, the animal was monitored for onset of apnea, defined as the animal’s chest ceasing to rise and fall. If and when apnea occurred, the rat was placed in left lateral recumbency. The left thoracic wall was then auscultated continuously with a stethoscope to identify cessation of heartbeat (CHB). Following CHB confirmation, video recording was stopped. The observation chamber was cleaned between trials. When CHB did not occur within 20 minutes of IP injection, animals were euthanized with an overdose of carbon dioxide gas using a gradual fill (30% chamber volume per minute) technique. These cases were considered unsuccessful euthanasias.

### Necropsy Examination

Following CHB, each animal was carefully removed from the chamber and positioned in dorsal recumbency for necropsy examination. The skin was incised along the midline and the injection site was identified in the abdominal wall musculature. The abdominal wall was incised and the intestines were reflected out of the abdominal cavity. Distribution of blue injectate and any misinjection into hollow viscera were noted. The liver was reflected cranially and any presence of dye within the biliary vessels caused by uptake of injectate from the peritoneal cavity and subsequent biliary excretion was noted. The GIT from the cardia to the descending colon was removed and any intestinal segments with dye-stained serosa were opened to confirm or rule out intraluminal misinjection. Misinjection was defined as the presence of blue injectate within hollow viscera or subcutaneous tissues, or staining the fur. For each rat, the serosal surfaces of the abdominal wall injection site, the caudate liver lobe, and transverse sections of at least three intestinal sections were examined histologically after formalin fixation for evidence of acute inflammation or swelling of mesothelial cells. Evaluation was performed by a single board-certified veterinary pathologist (CK), who was blinded to treatment group assignments.

### Off-Line Video Analysis

Videos of the HH and LL trials were analyzed for the incidence of writhing behavior by a single individual blinded to treatment (JR). Baseline recordings were analyzed in their entirety while post-injection videos were analyzed until the rat became ataxic. Writhing was defined as a contraction of the lateral abdominal walls to the extent where the abdomen became concave with concurrent extension of the pelvic limbs [17, 22].

### Statistical methods

All statistical analyses were performed using commercial software (GraphPad Prism v.6.03, GraphPad Software, Inc. La Jolla, California, USA and IBM SPSS Statistics 21, IBM, Armonk, NY, USA). Data were considered approximately normal if skewness and kurtosis were less than ± 1.5 and 3, respectively. Between-group comparisons were performed with a one-way ANOVA with a Games-Howell multiple comparisons test. Consistency of the euthanasia process was assessed with a coefficient of variation (CV) calculation. A p-value of < 0.05 was considered significant. Data are presented as mean ± SD.

## Results

Of 51 trials, 43 (84.3%) were successful IP injections and 8 (15.7%) were misinjections. Successful IP injection resulted in death in all PB groups: 34 (79.1%) were given IP PB and 9 were control animals. Successful deaths were distributed as follows: LL (n = 11), LH (n = 12), and HH (n = 11).

The fastest killing method from injection to CHB was the HH group (283.7 ± 38.0s), which was significantly faster than both the LL (485.8 ± 140.7s, p = 0.002) and LH (347.7 ± 72.0s, p = 0.039) groups (Fig. 2). Euthanasia in the LH group was also significantly faster than the LL group (p = 0.027).

**Fig 2:**
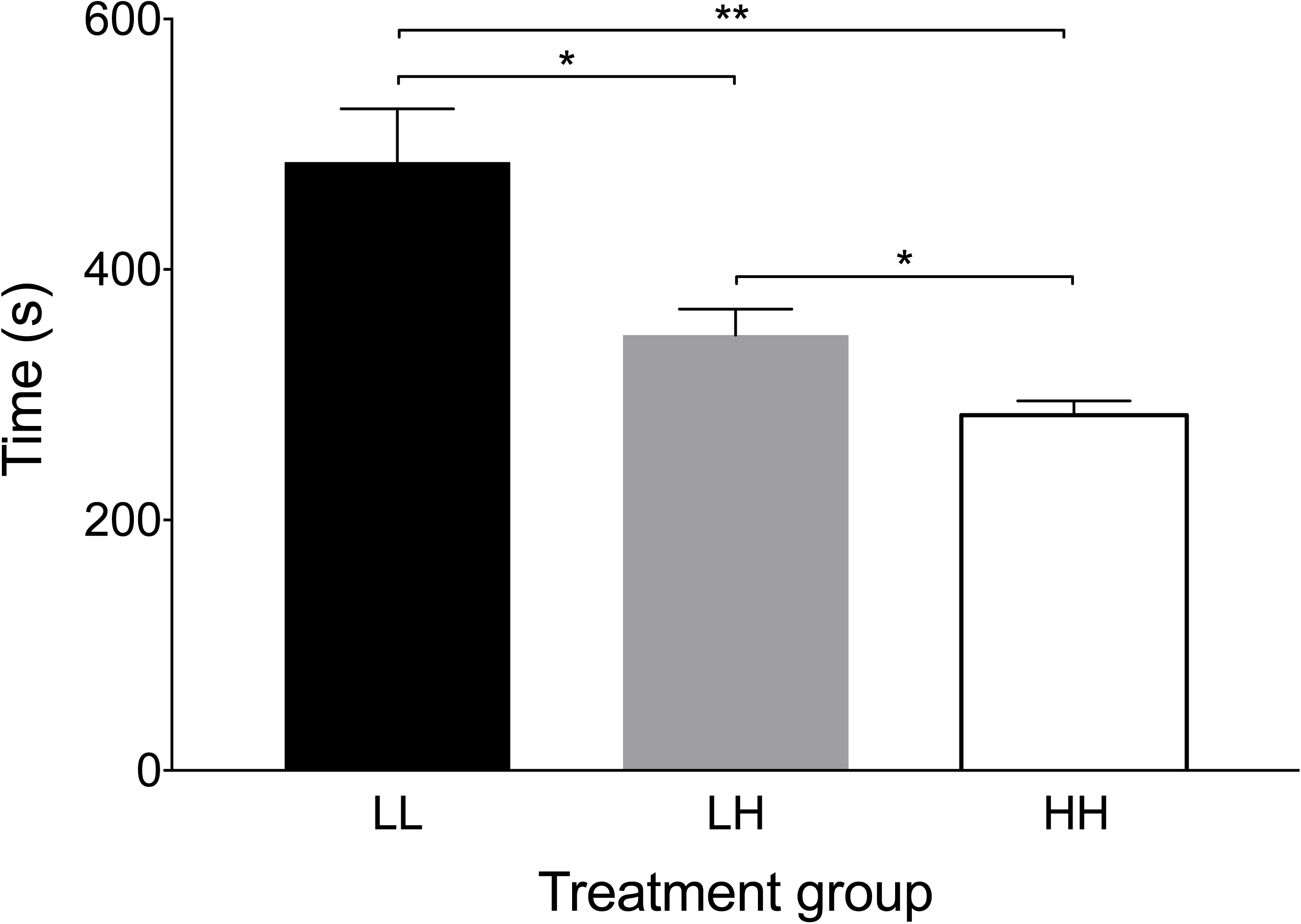
Time from delivery of the intraperitoneal injection to cessation of heart beat was fastest in the high-dose high-volume group (HH). LL = low-dose low-volume group, LH = low-dose high-volume group. *p < 0.05 **p = 0.002

The HH group was not only the fastest, but also the most consistent euthanasia method. The CV for HH was 13.4%, compared with 29.0% for LL and 20.7% for LH groups.

The period from injection to LORR was longest in LL (139.5 ± 29.6s), compared with both HH (111.6 ± 19.7s, p = 0.046) and LH (104.2 ± 19.3s, p = 0.01, Fig 3A). Time from injection to LORR did not differ between LH and HH (p = 0.64). The LORR-apnea time period showed the greatest variation between treatment groups and therefore had the greatest influence on the speed of the overall time to death (Fig 3B). LORR-apnea was significantly faster in the HH group (56.8 ± 25.1s) than LL (253.3 ± 106.7s, p < 0.001) and LH (146.6 ± 66.1s, p = 0.002). LORR-apnea in the LH group was also significantly faster than in the LL group (p = 0.03). There was no significant difference from apnea-CHB among treatment groups: HH (116.2 ± 19.7s) versus LH (93.0 ± 29.0s, p = 0.09), HH versus LL (92.9 ± 24.2s, p = 0.06), LH versus LL (p = 1.0).

**Fig 3:**
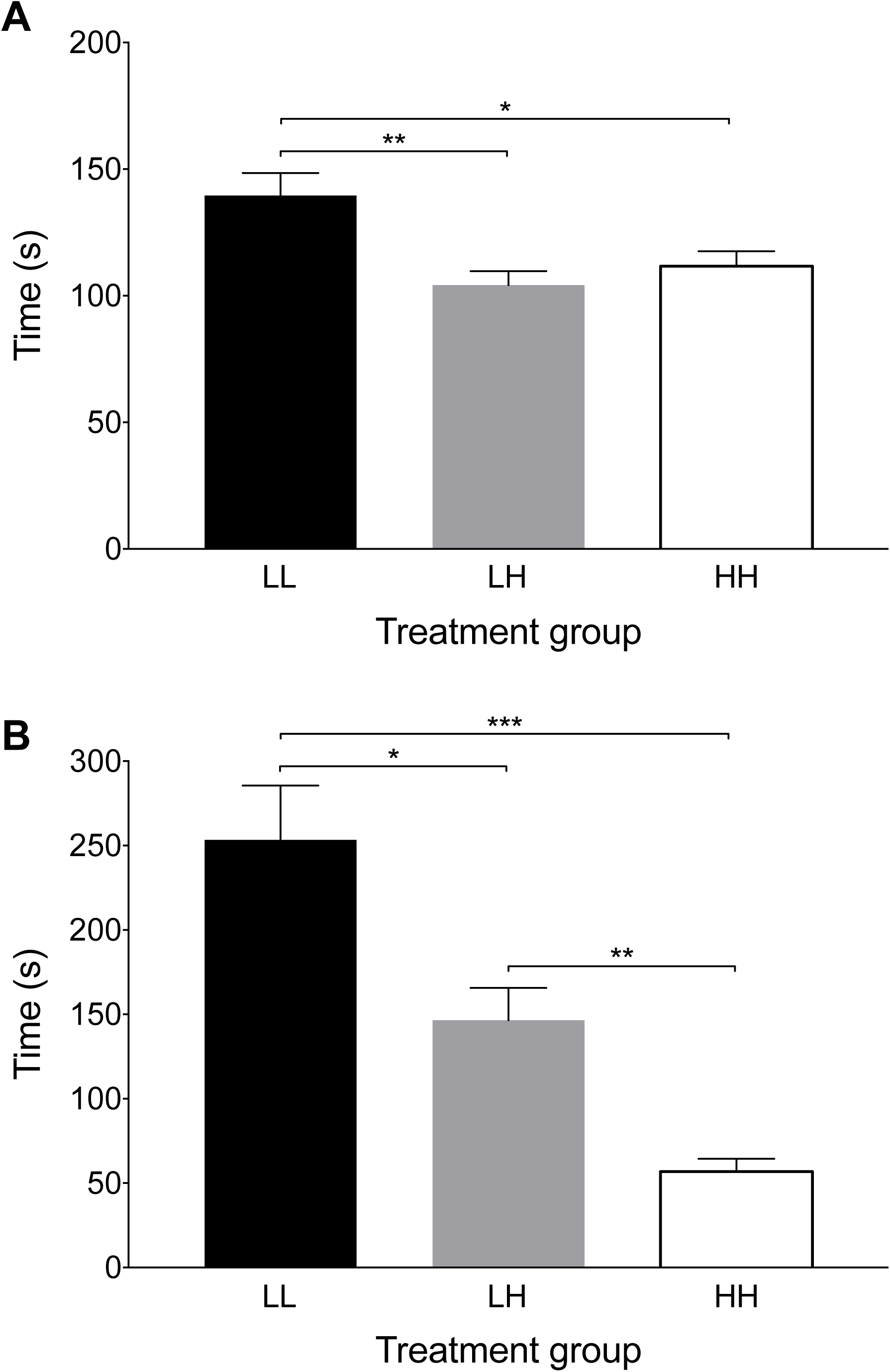
Time from delivery of the intraperitoneal injection to loss of the righting reflex was longest in the low-dose low-volume group (LL). LH = low-dose high-volume group, HH = high-dose high-volume group. *p < 0.05 **p = 0.01.Fig 3B: Time from loss of the righting reflex to apnea was shortest in the high-dose high-volume group (HH). *p < 0.05 **p = 0.002 ***p < 0.001

Eight misinjections were identified at necropsy. One misinjection was in a control animal. Seven misinjections were treatment group rats (HH; n = 3, LH; n = 2, LL; n = 2). Of these, euthanasia was unsuccessful (exceeding 20 minutes) in 3 (42.8%) animals (HH [n = 2], LL [n = 1]). In the four animals in which euthanasia was successful (HH [n = 1], LH [n = 2], LL [n = 1]), injection-CHB ranged from 318-1200s.

The anatomic distribution of the eight misinjections was as follows: four entered the cecal lumen (Fig 4B), two entered the jejunal lumen (Fig 4C), one was entirely within the subcutaneous tissues of the abdominal wall (Fig 4D), and one was predominantly over the fur of the medial thigh, with a small amount in the subcutaneous space. Cecal positions were variable: 14/51 (27.5%) in the right caudal quadrant, 5/51 (9.8%) located in the midline and 32/51 (62.7%) in the left caudal quadrant.

**Fig 4:**
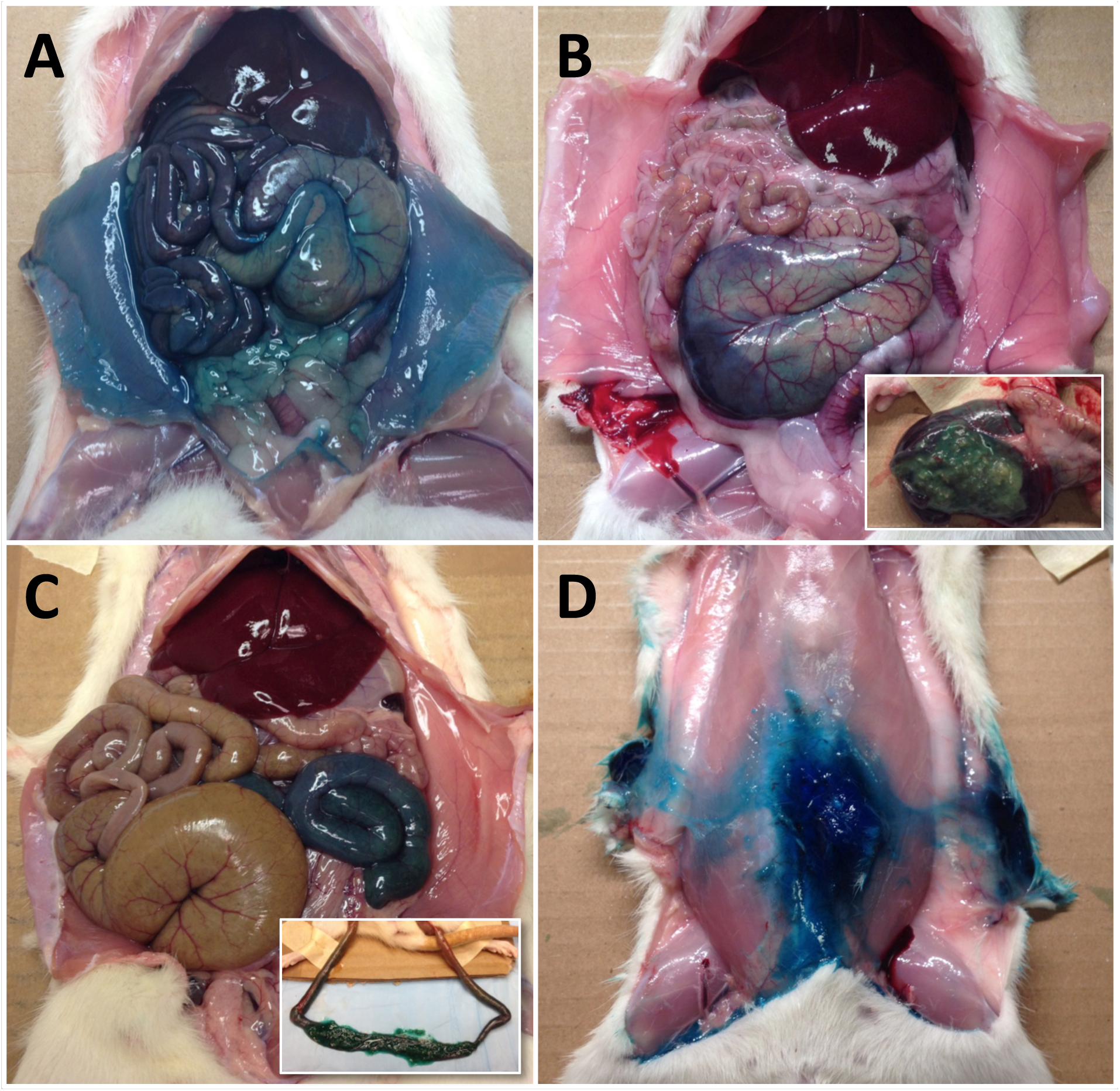
Abdominal cavities of four rats after confirmation of death; ventral view. Diffuse blue dye staining of serosal surfaces following successful intraperitoneal injection (A). Restricted dye distribution following inadvertent cecal (B), intestinal (C), and subcutaneous (D) misinjection. The insets in panels B and C show dye-stained ingesta, confirming inadvertent luminal misinjection.

### Writhing

Writhing was not observed in either the LL or HH groups in baseline video recordings. Following injection, writhing, assessed in animals with successful injections, was seen in 45.5% (5/11) of HH and 36.4% (4/11) of LL rats.

## Discussion

Historically, concerns regarding the variable success of IP euthanasia have revolved around misinjection leading to variability and potential pain [7, 16–20, 22].

Our results show that: 1. IP injection with 800 mg/kg sodium pentobarbital (HH group) resulted in the fastest and most consistent killing method; 2. variable cecal position contributed to misinjections; and 3. the incidence of writhing behavior was less than half of that previously reported.

Both dose and volume contribute to the speed of euthanasia, and dose in particular appears to have the most dramatic effect on consistency of technique. The speed and consistency of the killing process can be improved through an increase in dose (accompanied by an increase in volume). Increasing injectate volume without increasing dose (LH group) improved the speed of IP euthanasia. However, further improvements in speed and consistency were achieved in the HH group.

From these results, several conclusions can be drawn. The mean + 2SD for the period from completion of injection to LORR was 151.0 seconds when 800 mg/kg pentobarbital (HH group) is administered IP. Therefore, it is highly likely that an animal that maintains LORR beyond this time has experienced a misinjection. If using the period from completion of injection to apnea as the indicator of successful injection, the time for mean + 2 SD was 259.1 seconds. Should these times be exceeded, a second injection of pentobarbital or alternative killing method should be performed.

Any increase in pentobarbital use is associated with an increased cost. For the formulation used here, this equates to approximately US$0.13 for the HH technique in a 250g rat. While cost is an important consideration, it should be weighed against the labor cost of the slower (approximately 1.7 fold) LL group and potentially prolonged pain experience during the period from injection to LORR.

Misinjection is a consistent limitation of IP PB. The rate of misinjection in this study was consistent with the range reported in the literature (for injections given in to the caudal right abdominal quadrant), from 6 to 20% [18–20]. A factor contributing to the misinjection rate is variability in cecal position. IP injection is usually performed in the right caudal abdominal quadrant and previous work has confirmed that the cecum is most commonly located in the left caudal abdominal quadrant (61.9%, right 24.2%, middle 13.8%, total n = 289 adult male and female rats) [19]. Our results are similar to these findings despite using a different injection technique. In the study of Coria-Avila et al. (2007), rats were restrained by a single person and suspended vertically by the thorax with the head up. This suggests that body position during injection has a minimal effect on the incidence of misinjection. Based on the misinjection rates in this and other studies, as well as the positional variation of abdominal viscera noted on necropsy, any suggested method of IP euthanasia in unlikely to prevent completely the possibility of misinjections.

Given this inherent obstacle in refining the euthanasia process, we hope that the recommendations described above will facilitate early identification of a misinjection, guiding the decision to repeat the injection or select an alternative euthanasia method. We observed a substantially lower incidence of writhing behavior than previously reported [16, 17]. To facilitate comparison, we used the same definition of writhing as that described by Wadham (1996) and Ambrose (1998, 1999) [16, 17, 22]. The reason for this discrepancy is unclear and may result from several factors.

The proposed cause of writhing behavior is the pain resulting from the alkaline pH of the PB solution. The pH of the solution studied here was 11.02 (measured independently by a commercial compounding pharmacy) and that of Wadham (1996) and Ambrose (1998) ranged between 10.9-12.6 [16, 22]. Current suggestions to alter the effect of pH focus on changing solution pH through buffering or the addition of lidocaine to provide analgesia [3, 4]. Wadham (1996) reported that buffering a solution of sodium pentobarbital from an original pH of 12.6 to 9.4 resulted in precipitation [22].

Any study combining behavioral observation in the presence of drugs with sedative properties is inherently confounded by a reduced ability to express behaviors as sedation occurs. This is a limitation of the study design. The use of a vehicle control would address this, but one was not readily available as there were restrictions in obtaining formulation information from the manufacturer of PB. Furthermore, the dose we used in the LL group (200 mg/kg), was higher than that of Ambrose (1998, 1999) (150 mg/kg) and selected based on our institutional SOP [16, 17]. This may have contributed to the lower incidence of writhing observed by shortening the time after injection when writhing behavior could be expressed, before sedation occurred. Finally, a lack of habituation to handling may have contributed to our findings. The rats used in this study received little or no handling prior to the experiment. Therefore, the stress associated with handling, injection and the observation chamber may have led to a suppression of normal behaviors.

By coupling the effects of volume and dose with the incidence of misinjections we have suggested practical guidelines to refine overdose with IP sodium pentobarbital as a killing method in rats.

## Declarations

Ethics approval and consent to participate

The animal care and use protocol was approved by the Health Sciences Animal Care Committee of the University of Calgary (AC11-0044), in accordance with the guidelines of the Canadian Council on Animal Care.

### Consent for publication

Not applicable

### Availability of data and materials

The datasets generated and analysed during the current study are available in the Harvard Dataverse repository: Pang, Daniel, 2016, "rat IP PB dataset", <u>doi:10.7910/DVN/PMGCHG</u>, Harvard Dataverse, V1 [UNF:6:A8TqIyFIJXya5kXJ5eW0kQ==]

### Authors’ contributions

KZ contributed to study design, collected and analyzed data and wrote the first draft of the paper. CGK collected and analyzed data and revised the manuscript. JR analyzed data and revised the manuscript. DSJP conceived the study, contributed to study design, collected and analyzed data and revised the manuscript. All authors read and approved the final manuscript.

## Acknowledgements

Not applicable.

